# Internal microbial zonation assists in the massive growth of marimo, a lake ball of *Aegagropila linnaei* in Lake Akan

**DOI:** 10.1101/2021.03.20.434239

**Authors:** Ryosuke Nakai, Isamu Wakana, Hironori Niki

## Abstract

Marimo (lake ball) is an uncommon ball-like aggregation of the green alga, *Aegagropila linnaei.* Although *A. linnaei* is broadly distributed in fresh and brackish waters in the northern hemisphere, marimo colonies are found only in particular habitats. The colonies have been gradually shrinking in recent years. Nevertheless, it is not clear how and why *A. linnaei* forms such massive spherical aggregations. Here, we report the bacterial microbiomes inside various sizes and aggregating structures of natural marimo collected from Lake Akan, Japan. We observed multi-layers composed of sediment particles only in the sizeable radial-type marimo with a >20 cm diameter, not in the tangled-type marimo. The deeper layers were enriched by *Nitrospira*, potential novel sulphur-oxidizing bacteria, and sulphate-reducing *Desulfobacteraceae* bacteria. The sulphur cycle-related bacteria are unique to Lake Akan due to sulphur deposits from the nearby volcanic mountains. Some of them were also recovered from lake sediments. Microorganisms of the multi-layers would form biofilms incorporating nearby sediment, which would function as microbial “seals” within large radial-type marimo. We propose that the layer structure provides habitats for diverse bacterial communities, promotes airtightness of the marimo, and finally contributes to the massive growth of the aggregation. These findings provide a clue to deciphering the massive growth of endangered marimo aggregates.

## Introduction

*Aegagropila linnaei,* a filamentous green alga, widely inhabits ponds, lakes, and brackish waters at high latitudes in the northern hemisphere (Boedeker et al., 2010b). Morphologically, individuals of this species are filaments of 1–4 cm that are generally found attached to rocky substrates with rhizoids. This alga is known for an intriguing phenomenon in some lakes, including Lake Akan in Japan: it aggregates in spherical formations known as “lake balls” or “marimo” (Japanese for algal *ball*) (Soejima et al., 2009; Umekawa et al., 2021).

There are two structurally different types of marimo: the “tangled type” is an aggregate of disorderly entangled filaments, and the “radial type” is an aggregate of filaments radially arranged from the center to the surface (Horiguchi et al., 1998; Einarsson, 2014) (**Figure 1**). The size of the tangled-type is usually several cm in diameter and not over 15 cm. On the other hand, the radial type develops into a sizable ball that occasionally reaches about 30 cm in diameter through the tip growth of the filaments (Horiguchi et al., 1998). Undoubtedly, the environmental conditions in habitats are significant determinants for the formation of the tangled- and radial-type marimo. Indeed, each type inhabits different areas of Lake Akan with different environmental conditions (**Extended Data Figure 1**).

**Figure 1.**
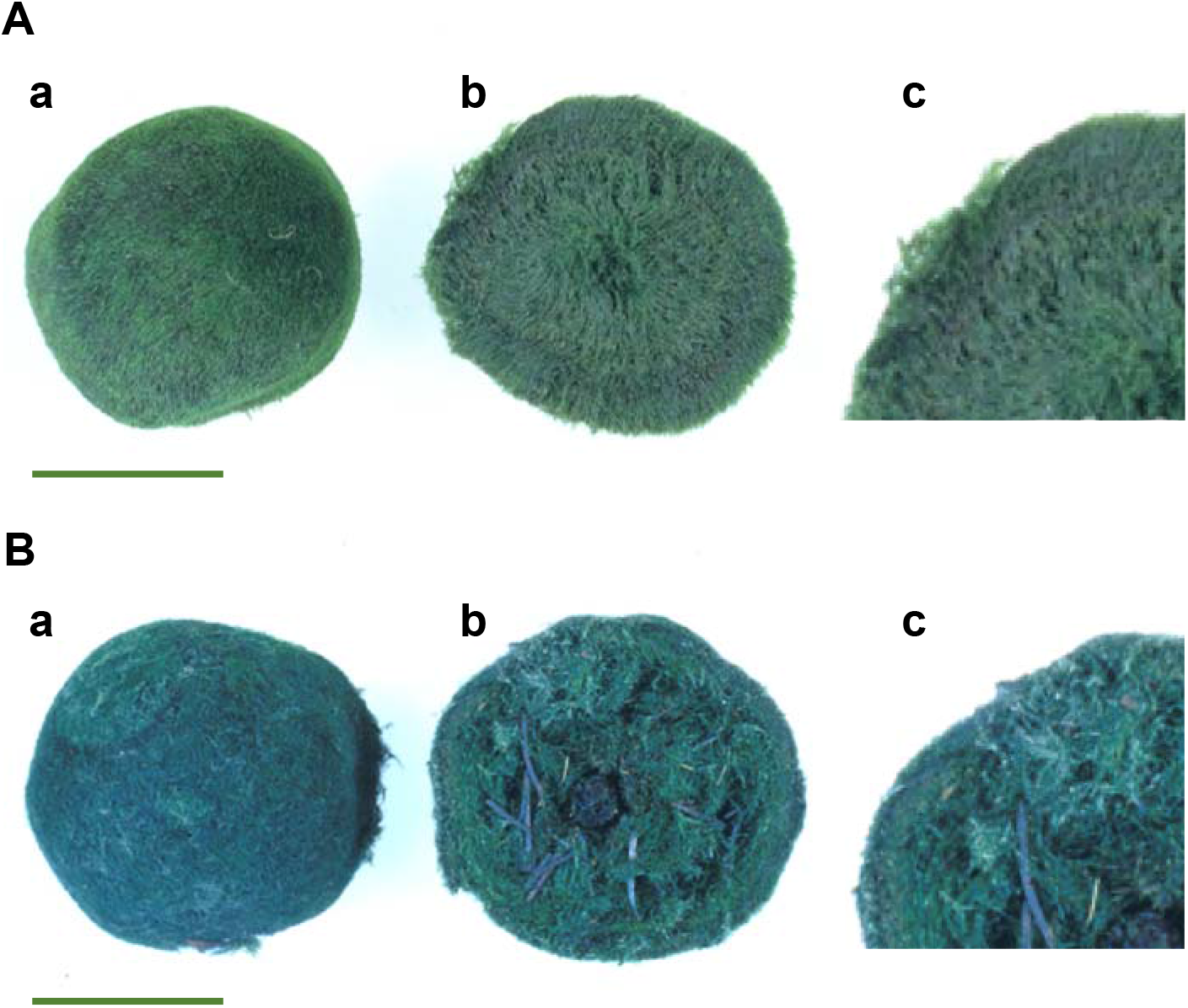
Two structurally different types of marimo from Lake Akan. (**A**) Images of the radial-type marimo: (a) outward appearance, (b) a cross section showing the radial arrangement of filaments, (c) an enlarged cross section. (**B**) Images of the tangled-type marimo: (a) outward appearance, (b) a cross section showing the entanglement of pine leaves and an alder cone in the tangled filaments, (c) an enlarged cross section. Scale bar, 5 cm.

Recently, dense colonies of the radial-type marimo that grow over 10 cm in diameter have been found in two lakes, Lake Mývatn, Iceland and Lake Akan (Einarsson et al., 2004), although this species inhabits many lakes. Regrettably, the marimo in Lake Mývatn have almost disappeared due to a deterioration of the environmental conditions (Einarsson, 2014). In Lake Akan, the dense colonies of the radial-type marimo are maintained within a restricted area at Churui Bay and Kinetanpe Bay.

It is known that external factors such as light, water quality, and water motion are involved in giving the marimo its spherical shape (Boedeker et al., 2010a; SANO et al., 2016). In addition to environmental conditions, we considered that various internal factors might be crucial for the more massive growth of the radial-type marimo. When the radial-type marimo inhabiting Lake Akan grows up about 10 cm in diameter, the filaments near its center begin to die, and a hollow is formed inside (Horiguchi et al., 1998). As a result, the densely packed filaments remain as an outer mass. These hollow structures are specific to the large aggregations of the radial-type marimo and are not formed in the smaller marimo of several cm in diameter. In a similar phenomenon, large aggregates of aquatic mosses are known to form “Koke-Bouzu,” or large pillar-like structures with an internal hollow, at the bottom of ultraoligotrophic Antarctic lakes (Imura et al., 1999). The formation of the hollow structure in Koke-Bouzu is positively related to the internal microbiomes (Nakai et al., 2012). The interior of Koke-Bouzu is decomposed and rotten; redox-potential gradients between this rotten interior and the exterior lead to a high diversity of microorganisms that are involved in material cycling in the *in situ* environment (Nakai et al., 2019). Therefore, we presumed that the hollow structure and its associated microbiomes are related to the large size of the radial-type marimo.

## Results and Discussion

We dissected three differently sized natural samples of the radial-type marimo, each having a different internal structure and different redox conditions. Then we examined the microbial communities in the outer and inner sections of these fresh marimo samples.

We collected the radial-type marimo at Churui Bay in Lake Akan, and categorized them into three sizes: small (about 4 cm in diameter, *n* = 3), medium (11 to 12 cm in diameter, *n* = 3), and large (21 and 22 cm in diameter, *n* = 2). Each of the samples was cut to separate the outer and inner sections of the algal outer mass on a board immediately after the collection (**Figure 2A**). Using 16S ribosomal RNA (rRNA) gene-based metabarcoding against pooled DNA samples of each size fraction, we constructed bacterial catalogs for the sections of the marimo of different three sizes, as well as for other samples, including the tangled-type marimo, the filaments form, and the surrounding lake water and sediments, for comparison **(Figure 2B and Supplementary Tables 4–9)**. We excluded archaeal taxa from further analyses because they were poorly represented in our metabarcoding data. We found no apparent differences in the non-parametric Shannon diversity index (*α* diversity) among the marimo microbiomes **(Supplementary Table 2**). This result indicates that the species richness of microbiomes is similar in the outer and inner sections. However, the *β* diversity calculated by a generalized UniFrac dendrogram indicated apparent similarities and differences among bacterial communities of the marimo microbiomes (**Figure 2C**). The UniFrac dendrogram showed that although both the microbiome in the outer and that in the inner section of the small-sized marimo were closely clustered, the microbiomes in the inner and outer sections of the medium- and large-sized marimo formed separate clusters. *Nitrospira* was frequently detected in the inner sections, while cyanobacteria were mainly found in outer sections in the larger radial-type marimo **(Figure 2B)**. In particular, filamentous cyanobacterial members of *Nostocales* and *Oscillatoriales* were dominantly detected (**Supplementary Table 6**). Filamentous cyanobacteria are generally considered to form benthic mats in aquatic environments, and their nitrogen-fixing capabilities are nutritionally significant in the biotic community (Paerl et al., 2000). Although the water quality of Lake Akan is classified as mesotrophic, it is known to become nitrogen deficient during the summer season, when the production and growth of aquatic plants and phytoplankton are active. Our measurements in Churui Bay, where we collected the marimo, showed that the concentration of PO_4_^3−^ as well as the concentrations of NH_4_^+^, NO_2_^−^, and NO_3_^−^ in the surface water were below the detection limits (**Supplementary Table 3**). Therefore, cyanobacterial N_2_ fixation might help in the growth of marimo under a nitrogen-deficient condition.

**Figure 2.**
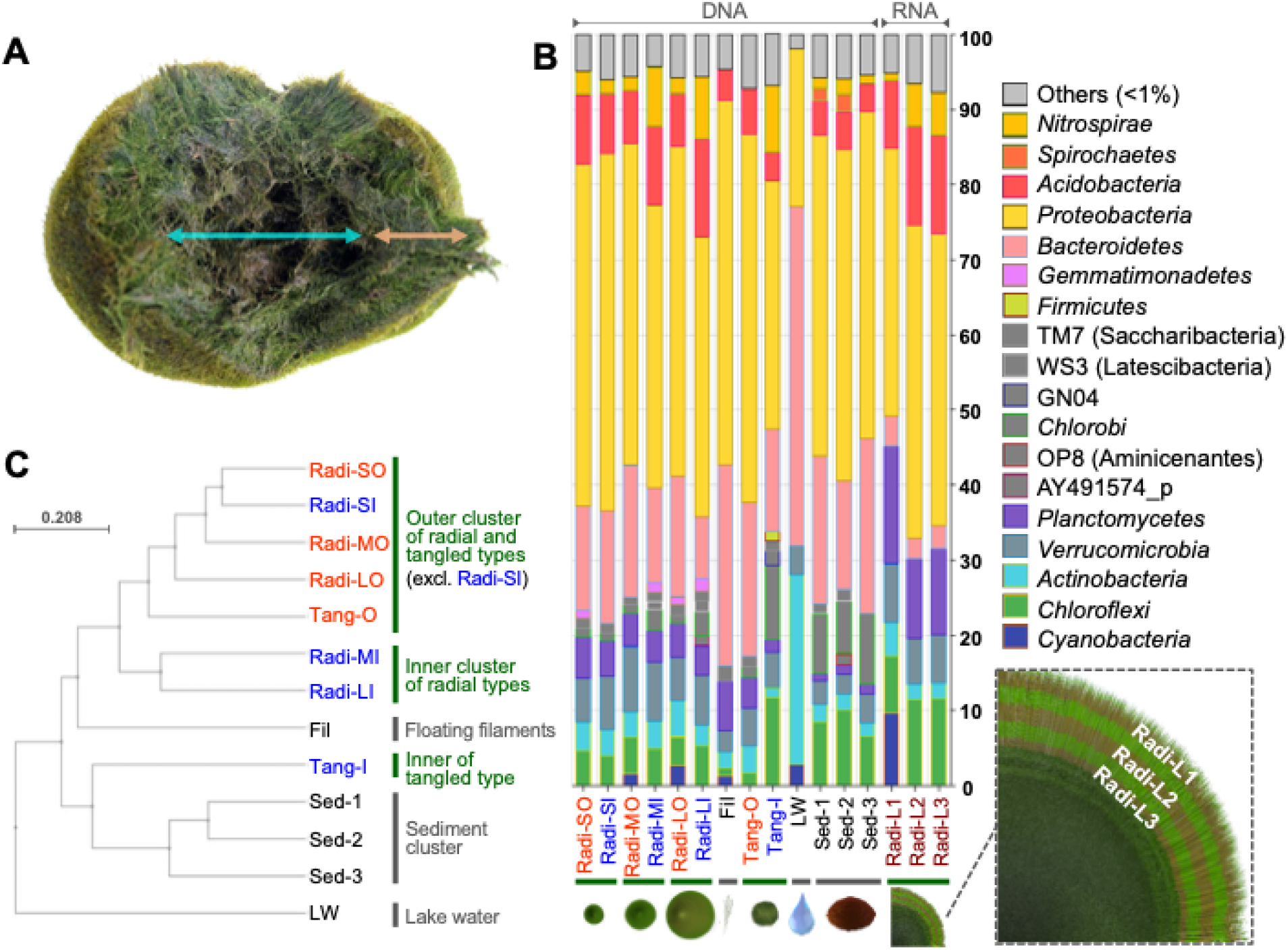
An integrated view of the phylogenetic diversity of the internal microbiomes of natural marimo. **(A)** Images of the interior structure of the radial-type marimo with a major axi of about 12 cm. The algal outer mass is indicated by an *orange arrow*, and the hollow structure is indicated by a *blue arrow*. **(B)** Taxonomic composition of 13 bacterial phyla and 5 candidate phylum-level lineages (TM7, WS3, GN04, OP8, and AY491574_p) with >1% sequence abundance in at least one sample in DNA-based 16S rRNA gene amplicon data from radial-type marimo, floating filaments, tangled-type marimo, and other environmental samples, and RNA-based 16S rRNA transcript amplicon data from the multi-layers that developed only in the large radial-type marimo samples (see the layer structure presented in **Figure 3**). The outer sections of small-, medium-, and large-sized radial-type marimo are labeled Radi-SO, Radi-MO, and Radi-LO, respectively, and the inner sections are labeled Radi-SI, Radi-MI, and Radi-LI, respectively. Floating *A. linnaei* filaments and the outer and inner sections of the tangled-type marimo were labeled Fil, Tang-O, and Tang-I, respectively. Surface lake water and sediments surrounding the studied marimo colony were labeled LW, Sed-1, Sed-2, and Sed-3. The multi-layers (three layers), which were composed of sediment particles, were labeled Radi-L1, Radi-L2, and Radi-L3. Details for each sample are given in **Supplementary Table 1**. **(C)** A generalized UniFrac dendrogram showing similarities among the bacterial microbiomes.

Meanwhile, nitrite-oxidizing *Nitrospira* members also can convert urea to NH_4_^+^ and CO_2_ and are considered to contribute to nitrogen (N) cycling processes beyond nitrite oxidation (Koch et al., 2015). Thus, taxonomically different but functionally relevant (i.e., N cycle-related) bacteria were characteristically recovered from the outer and inner sections of the larger marimo. In contrast, such a distribution was not found in the smaller marimo. The UniFrac dendrogram also indicates that the marimo microbiomes were separately clustered with the other samples, including the surrounding lake water and sediments. These results suggest that the radial-type marimo harbor unique and specific bacterial communities that assist in massive growth regardless of poor nutrient conditions.

We analyzed the interior structure of the radial-type marimo with a diameter of >20 cm. A cross-section of the marimo showed that the algal outer mass developed to a thickness of about 2.5 cm, and the core part had a hollow. Additionally, we found multi-layers composed of accumulated brown-colored sediment particles inside the outer mass. Note that the sizable marimo populations distributed in shallow areas tend to be covered by lake sediments (**Extended Data Figure 2**), while the attached sediments appear to be removed by the marimo’s rotation. However, the sediment particles themselves are often sandwiched in layers within the algal outer mass. The large marimo we collected had a three-layered structure, and we named the layers Radi-L1 (0.3 to 0.4 cm deep from the surface), Radi-L2 (1 cm deep), and Radi-L3 (2 cm deep) (**Figure 3A and Extended Data Figure 3**). We then determined the composition of viable microbiomes in these multi-layers by means of RNA-based metabarcoding of the 16S rRNA transcripts. We used RNA for this purpose because 16S rRNA transcripts are mainly isolated from living bacterial cells, but not dead cells. The results of the RNA-based metabarcoding analysis showed that the dominant bacterial groups were common to all samples, as shown in DNA-based metabarcoding analyses (**Figure 2B**). However, distinctive distributions of certain taxa were found in each layer. Some of the dominant bacterial genera and genus-level lineages were different between the surface layer, Radi-L1, and the deeper layers, Radi-L2 and Radi-L3 (**Figure 3B**). In Radi-L1, a novel cyanobacterial lineage within the *Symploca*-related family was frequently detected. The novel δ-proteobacterial lineage AJ617907_g of *Polyangiaceae* and *α*-proteobacterial *Hyphomicrobium* were also detected. In contrast, these members were minor components (<0.2%) in the Radi-L2 and Radi-L3 layers. *Symploca* spp. are aerobic, filamentous N_2_-fixing cyanobacteria that occur in microbial mats (Stal, 2012). The δ-proteobacterial AJ617907 sequence was initially discovered from the oxic-anoxic interphase of soil. Members of *Hyphomicrobium* are considered to be significant players in denitrification (Martineau et al., 2015), which is a microbial process of converting NO_2_^−^ and NO_3_^−^ to N_2_ under oxygen-limiting conditions. The detection of these viable members suggests that an O_2_ concentration gradient might be formed in the surface layer Radi-L1.

**Figure 3.**
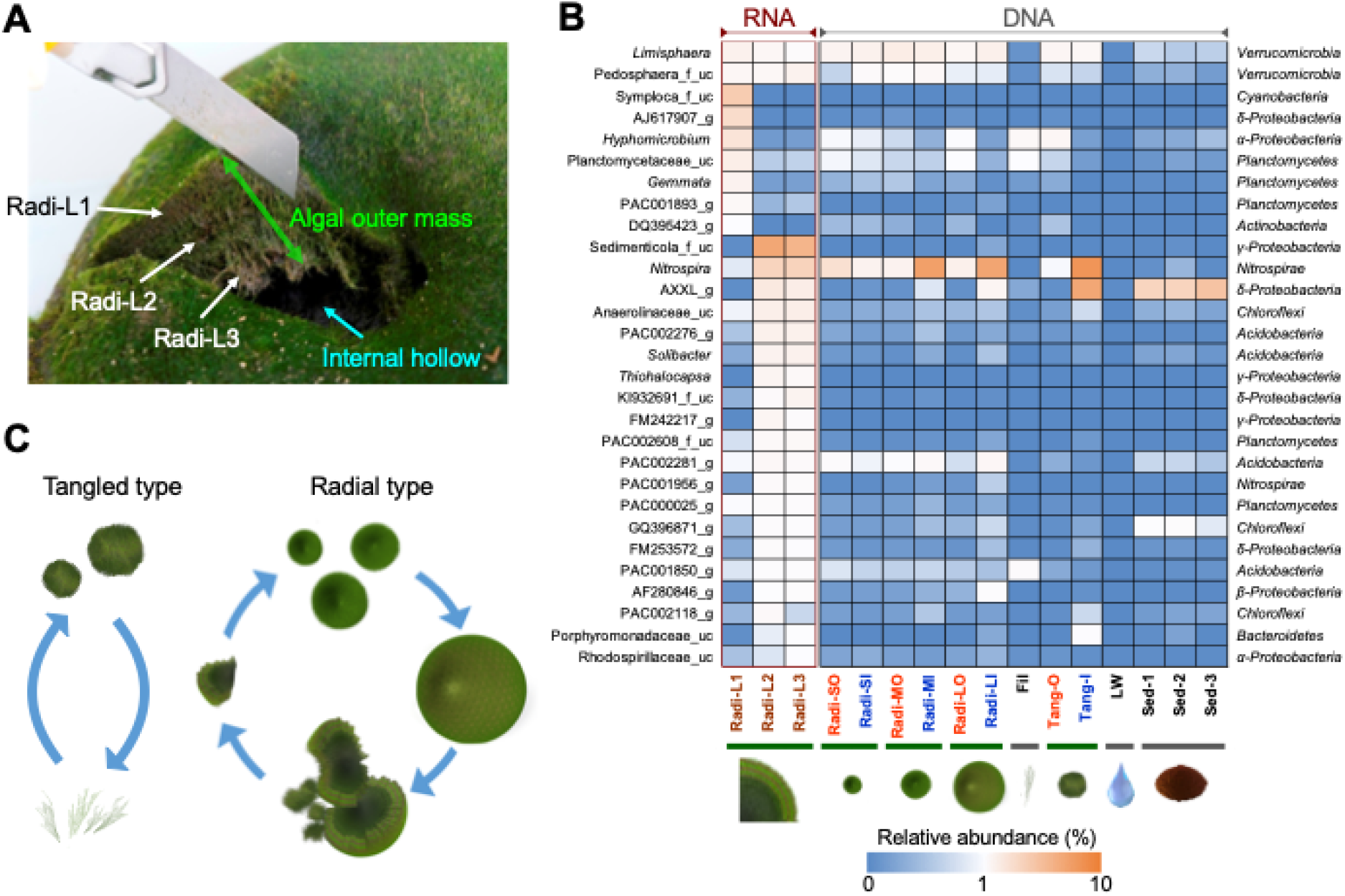
Multi-layers, internal microbial zonation, and proposed development process of the sizable radial-type marimo. (**A**) Image of three brown-coloured layers inside the algal outer mass and the internal hollow of the large-sized, radial-type marimo with a diameter of >20 cm. The three layers were labeled Radi-L1 (0.3 to 0.4 cm deep from the surface), Radi-L2 (1 cm deep) and Radi-L3 (2 cm deep). (**B**) Heatmap showing six genera (*Limisphaera*, *Hyphomicrobium*, *Gemmata*, *Nitrospira*, *Solibacter*, and *Thiohalocapsa*) and 23 other unclassified or candidate genus-level lineages with >1% sequence abundance in at least one sample in RNA-based 16S rRNA transcript amplicon data from the multi-layers. The frequency of occurrence of each lineage in other samples obtained from DNA-based 16S rRNA gene amplicon data is also shown (note that sample name codes are presented in **Figure 2** and **Supplementary Table 1**). EzTaxon category names and taxonomic affiliations at the phylum or class level (for *Proteobacteria*) are shown on the *left* and *right side*, respectively. Following the EzBioCloud database, the uncultured phylotype is tentatively given the hierarchical name assigned to the DDBJ/ENA/GenBank accession number with the following suffixes: “_s” (for species), “_g” (genus), “_f” (family), “_o” (order), “_c” (class) and “_p” (phylum). The unclassified sequences below the cut-off values are labeled “_uc” (for unclassified). (**C**) Models of development and regeneration of the tangled-type and radial-type marimo. For the tangled-type marimo, *A. linnaei* filaments cycle between a free-floating state and an entangled state (*left*). The radial-type marimo is regenerated from a fragment derived from a broken larger radial-type individual, so that the radial arrangements of filaments and the inside microbiomes are inherited (*right*).

On the other hand, a novel γ-proteobacterial group within the *Sedimenticola*-related family, the δ-proteobacterial AXXL_g of *Desulfobacteraceae*, *Nitrospira*, and other multiple uncultured lineages dominated in both Radi-L2 and Radi-L3 (**Figure 3B**). The novel γ-proteobacterial group and *Nitrospira* were especially highly enriched in Radi-L2 and Radi-L3 (6.1–7.1% and 3.9–4.0%, respectively), and were minor in Radi-L1 (0.03% and 0.8%, respectively). Known *Sedimenticola* strains are chemoautotrophic bacteria and can utilize sulphur compounds (e.g., elemental sulphur and hydrogen sulphide) (Flood et al., 2015). They fall within the unclassified *Gammaproteobacteria* monophyletic clade with multiple microaerobic sulphur-oxidizing endosymbionts of marine invertebrates (e.g., bivalve mollusc) endemic to sulfidic environments (Dubilier et al., 2008). The majority of the sequences of the novel γ-proteobacterial group had 93–94% similarity with the marine sediment sequence (DDBJ/ENA/GenBank accession no. JF344428). Their closest type strain was *Sulfurivermis fontis* JG42^T^, which is a sulphur-oxidizing autotroph that was isolated from a hot spring microbial mat (Kojima et al., 2017). The novel member inhabiting the marimo deeper layers might have a sulphur-related chemoautotrophic metabolism.

Filaments in the inner sections maintain their green color, indicating that chloroplasts remain despite the darkness within the marimo **(Figure 3A)**. When the inner filaments are isolated and irradiated with light, the photosynthesis activity recovers within a few days (Yokohama, 1994; Horiguchi et al., 1998). Thus, the photosynthetic activity of the radial-type marimo decreases from the surface to the interior, but the inner filaments are somehow viable. In this context, potential chemoautotrophic bacteria identified in the inner sections may carry out chemosynthetic CO_2_ fixation. Then the primary production by the CO_2_ fixation may contribute to the survival of inner filaments in the marimo interior. Remarkably, Lake Akan is rich in sulphur and hydrogen sulphide, which sulfur-oxidizing bacteria can use, due to the inflow of water from a river associated with a nearby volcano, Mt. Me-Akan (Suzuki et al., 1957; Japan Wild life Research Center, 2015).

Moreover, an uncultured lineage AXXL_g dominantly detected in the deeper layers of Radi-L2 and Radi-L3 belongs to *Desulfobacteraceae* within *Deltaproteobacteria*. Most of the *Desulfobacteraceae* species are sulphate-reducing anaerobes (Kuever, 2014). The AXXL_g-related sequences were found in the sediments surrounding the marimo colonies (**Figure 3B**). Other lineages, such as GQ396871_g of *Chloroflexi* in Radi-L2 and Radi-L3, were also enriched in the nearby sediments. The spatial distributions suggested that the two deeper layers possess anaerobic microenvironments in which sediment bacteria can inhabit.

It is noted that the filamentous cyanobacteria on the marimo surface might be involve in not only CO_2_ and N_2_ fixation but also the incorporation of minerals and organic matter. Furthermore, the filamentous microorganisms produce extracellular polymeric substances and them release into their surroundings (Stal, 2012). The surfaces of *A. linnaei* filaments isolated from the radial-type marimo are known to be highly sticky. Cyanobacteria-mediated formation of microbial granules and of biofilms is often observed in nutrient-limited glaciers and polar freshwater lakes (Uetake et al., 2016; Jungblut and Vincent, 2017). In addition, many of the *Acidobacteria* that were frequently detected in both Radi-L2 and Radi-L3 (**Figure 3B**) have the potential to form biofilms and facilitate particle-aggregates (Kielak et al., 2016). Thus, it is possible that “microbial sealing” by bacterial aggregates/biofilms exerts beneficial effects on the strength of the multi-layers inside marimo. In fact, sizeable radial-type marimo are highly airtight and rigid structures (**Extended Data Figure 4 and Supplementary Movie 1**). It is difficult for water to enter the internal cavity. Once the inside water is removed, the marimo floats on water for days (**Extended Data Figure 4C**).

Physical factors in the growing environment are involved in the developmental mechanism of the ball-like marimo. Wind waves are a driving force which rotates marimo and maintains their rounded spherical form (SANO et al., 2016). The rotation of marimo enables photosynthesis over the entire marimo-surface and removes sediment from the surface. However, the tangled-type marimo cannot grow into a larger aggregate of over 20 cm in diameter like the radial-type marimo. Only the actively growing marimo over 20 cm in diameter developed the internal multi-layers. The layers were available for habitats of distinctive bacteria. As a result, the bacterial zonation would supply carbon sources and nutrients to the host *A. linnaei* and help the massive growth of the radial-type marimo. In addition, the microbial sealing of the layer structure could confer mechanical strength to the radial-type marimo. A possible scenario of the regeneration of a radial-type marimo is as follows **(Figure 3C)**. First, an extremely large marimo is broken down and fragmented by external forces containing water flow (**Extended Data Figure 2A**). Fragments of the large aggregate then reform as a small marimo due to inherit the radial arrangements of filaments and the interior microbiomes. Environmental eutrophication affects not only the algal growth, but also the microbial community inside the marimo. In combination with external factors, the internal microbiomes of the sizeable radial marimo provide clues that could help in the management of globally-endangered marimo colonies and the regeneration of massive aggregates in their former habitats.

## Supporting information

Supplementary Movie 1.mov

## Acknowledgments

We are deeply grateful to Drs. S. Imura and M. Tsujimoto for their invaluable help with the field sampling and diving. This work was partly supported by the Systematic Analysis for Global Environmental Change and Life on Earth (SAGE) project promoted by the Trans-disciplinary Research Integration Center, under the umbrella of the Research Organization of Information and Systems, Japan. This work was partly supported by a Grant-in-Aid for Scientific Research from the Japan Society for the Promotion of Science (no. JP15H05620 to RN).

## Author contributions

RN and HN coordinated the study. All authors conducted field studies. RN conducted microbiome experiments and data analysis. WI has observed the life cycles of marimo for many years and took all our underwater photos of the marimo in Lake Akan by scuba diving. All authors discussed the data, wrote the manuscript, and approved the final version.

