## Supplementary material for "Internal microbial zonation assists in the massive growth of marimo, a lake ball of *Aegagropila linnaei* in Lake Akan": Methods, and Extended Data Figures 1-4, and Supplementary tables 1-3

### Table of Contents


Materials and Methods ..................... 2

Extended Data Figures ..................... 7
  Figure 1 7
  Figure 2 8
  Figure 3 9

Supplementary Tables ..................... 10
  Table 1 10
  Table 2 11
  Table 3 12
  A note on Tables 4 to 9 13

Supplementary Movie ..................... 13
  A note on Movie 1 13

### Methods

#### *Sample collection*

Radial-type marimo samples of *Aegagropila linnaei* were harvested on 5 August 2014 at Churui Bay in Lake Akan (43° 27′ N 144° 06′ E), Hokkaido, Japan. Small (about 4 cm in diameter, $n = 3$), medium (11 to 12 cm, $n = 3$), and large (21 and 22 cm, $n = 2$) samples were collected by scuba divers. Each sample was immediately transferred to a researcher in a boat. The samples were dissected on the boat; the outer surface and inner sections (~1.5 cm × ~1.5 cm × ~0.5 cm portions) used for DNA extraction were picked apart using sterilized tweezers and were separately placed in sterilized tubes; the samples for RNA extraction were also picked apart and immersed in RNA*later*® (Thermo Fisher Scientific, Waltham, MA). The outer sections of the small, medium, and large samples were labelled Radi-SO, Radi-MO, and Radi-LO, respectively; the inner sections were labelled Radi-SI, Radi-MI, and Radi-LI. All samples were transported to the lakeside laboratory and stored at –80°C until analysis. The samples in RNA*later*® were kept at 4°C overnight and then deep-frozen. To characterise microorganisms associated with radial-type marimo, floating *A. linnaei* filaments (labelled Fil) and tangled-type marimo of about 5 cm in diameter ($n = 3$) were collected for comparative analysis at Takiguchi Bay in Lake Akan. The tangled-type samples were dissected into the outer surface and inner sections similarly to the radial-type marimo, and were then labelled Tang-O and Tang-I, respectively. Additionally, surface lake water (labelled LW) and three samples of sediment (labelled Sed-1, Sed-2, and Sed-3) surrounding marimo colonies were collected. These samples were stored as described above. Further details of the sample codes are presented in **Supplementary Table 1**.

#### *Evaluation of lake water quality*

The pH and electrical conductivity of the lake surface water were measured using a pH meter (D-71; Horiba, Kyoto, Japan) and a conductivity meter (ES-51; Horiba), respectively. The concentrations of $Na^+$, $K^+$, $Ca^{2+}$, and $Mg^{2+}$ were determined using an atomic absorption spectrometer (SOLAAR S Series; Thermo Fisher Scientific). The concentration of $NH_4^+$ was measured by indophenol blue absorptiometry (UV–160; Shimadzu, Kyoto, Japan). The concentrations of $Cl^-$, $NO_2^-$, $NO_3^-$, $SO_4^{2-}$, and $PO_4^{3-}$ were determined by ion chromatography (761 Compact IC; Metrohm, Herisau, Switzerland). To determine the total nitrogen concentration, the water sample was digested with potassium persulfate (Solórzano and Sharp, 2020) and was analysed by UV-spectrophotometry (UV–160; Shimadzu). Total phosphorus was determined by standard molybdenum-blue colorimetry following persulfate digestion (Menzel and Corwin, 1965). Total organic carbon was measured using a TOC Analyzer (TOC-VCPH; Shimadzu).

#### *DNA extraction, PCR amplification, and high-throughput sequencing*

DNA was extracted from natural marimo samples using a modified version of the bead-beating method described previously (Nakai, R. et al., 2012). Briefly, a 0.5-g sub-sample was placed in a Lysing Matrix E Tube (MP Biomedicals, Santa Ana, CA) and lysis solutions, pre-warmed at 65°C, of the ISOIL for Beads Beating Kit (Nippon Gene, Toyama, Japan) were added according to the manufacturer's manual. After shaking in a Beads Crusher μT-12 (Taitec, Saitama, Japan) at 3200 rpm for 45 s, the tube was incubated at 65°C for 1 h, followed by centrifugal separation. DNA was purified from the aqueous supernatant using a MagExtractor™ Genome Kit (Toyobo, Osaka, Japan). The above procedure was conducted three times for each sample of each size, and the extracted DNAs were pooled by each size group (that is, small, medium, or large) to create a representative DNA sample. PCR amplicons were generated using the universal primer set 342F-806R (Mori, H. et al., 2014) covering both archaeal and bacterial 16S rRNA genes. The PCR mixture composition, reaction conditions, and sequencing library generation followed the '16S Metagenomic Sequencing Library Preparation' protocol provided by Illumina (San Diego, CA, USA). Pair-end 300-bp MiSeq (Illumina) sequencing was performed using a Nextera XT Index Kit (Illumina). DNA from comparative samples was extracted as outlined above. For 2 l of the lake water sample, DNA extraction was carried out after filtering the sample through a 0.22-μm-pore Sterivex filter unit (Millipore, Billerica, MA) as described previously (Somerville et al., 1989). Each DNA sample was then PCR-amplified and sequenced as described above.

***RNA extraction and cDNA synthesis***

The internal multi-layers composed of sediment particles were observed only in the two large-sized, radial-type marimo samples. The three layers, the surface, intermediate and innermost layer were used from the surface to the inner were used for RNA extraction. The parts of each layer submersed in RNA*later*® were picked apart and labelled Radi-L1 (about 0.3 to 0.4 cm deep from the surface), Radi-L2 (1 cm deep), and Radi-L3 (2 cm deep), respectively. Total RNA was extracted from 0.5 g of each layer part using a FastRNA® Pro Soil-Direct Kit (Qbiogene, Carlsbad, CA), as per the manufacturer's instruction. Residual DNA was digested using a TURBO DNA-*free*™ Kit (Thermo Fisher Scientific). The absence of DNA contamination was confirmed as the absence of PCR amplification of the digested template using the universal primer set 342F-806R as indicated above. The RNA was reverse-transcribed into cDNA using a SuperScript® VILO cDNA Synthesis Kit and Master Mix (Thermo Fisher Scientific), including random primers. The above procedure was performed three times for each sample, and the cDNAs from each layer group were pooled. Each cDNA was PCR-amplified and sequenced as described in the "DNA extraction" section.

***Data analysis***

The MiSeq pair-end reads were assembled using fastq-join (Aronesty, 2013). After trimming the barcode and primer sequences, ambiguous sequences that were <80 nt and had a low average quality score (<25) were removed. Taxonomic assignment of the quality-filtered sequences was processed with the default settings in the Microbiome Taxonomic Profiling (MTP) pipeline of the EzBioCloud (https://www.ezbiocloud.net/contents/16smtp; ChunLab Inc., Seoul, Korea) (Yoon, S.-H. et al., 2017). This tool was previously called BIOiPLUG (ChunLab, Seoul, Korea). Briefly, the quality-filtered sequences were compared to the EzBioCloud 16S rRNA gene sequence database v. PKSSU4.0 constructed from the curated 16S rRNA gene sequences using the USEARCH program v.8.1.1861_i86linux32 (Edgar, 2020). It is noted that, in this database, the yet-uncultured phylotype is tentatively given the hierarchical name assigned to the DDBJ/ENA/GenBank accession number with the following suffixes: "_s" (for species), "_g" (genus), "_f" (family), "_o" (order), "_c" (class) and "_p" (phylum) (Kim, O.-S. et al., 2012). The taxonomic category was assigned based on the following cut-off values of sequence similarity: species (≥97%), genus ($>97\% > x \geq 94.5\%$), family ($>94.5\% > x \geq 86.5\%$), order ($>86.5\% > x \geq 82\%$), class ($>82\% > x \geq 78.5\%$), and phylum ($>78.5\% > x \geq 75\%$), where $x$ corresponds to the sequence identity with sequences in the database. These cut-off values were taken from previous studies (Yoon, S.-H. et al., 2017; Yoon, S.-H. et al. 2014). The sequences below the cut-off value at the species or higher level were assigned in the unclassified group labelled as "_uc" (for unclassified). Chimeras of the sequences that did not match at the species level (≥97%) were identified using the UCHIME program within USEARCH (Edgar et al. 2011) and the EzBioCloud chimera-free reference database (https://help.ezbiocloud.net/user-guide/mtp-pipeline/chimera-detection/), and chimeric sequences were omitted. Unmatched and eukaryotic plastid sequences were also excluded. The filtered valid sequences were used as the final dataset. Operational taxonomic units (OTUs) at a 97% identity threshold were picked up by the open-reference method (https://help.ezbiocloud.net/user-guide/mtp-pipeline/mtp-pipeline/) using the EzBioCloud MTP pipeline. Singleton sequences were removed in the OTU picking process according to Unno (2015). The sequence frequency within the classified taxa and the generalized UniFrac distance (Chen, J. et al., 2012) between samples were computed and visualized in the MTP pipeline. $\alpha$- and $\beta$-diversity indices were computed in the MTP pipeline.

***Nucleotide sequence accession number***

The MiSeq-derived sequence amplicon data ($n = 16$) are available at DDBJ/ENA/GenBank under the BioProject number PRJDB8727. The BioSample numbers are SAMD00224685 to SAMD00224700. The DRA accession number is DRA010176.

### Extended Data Figures

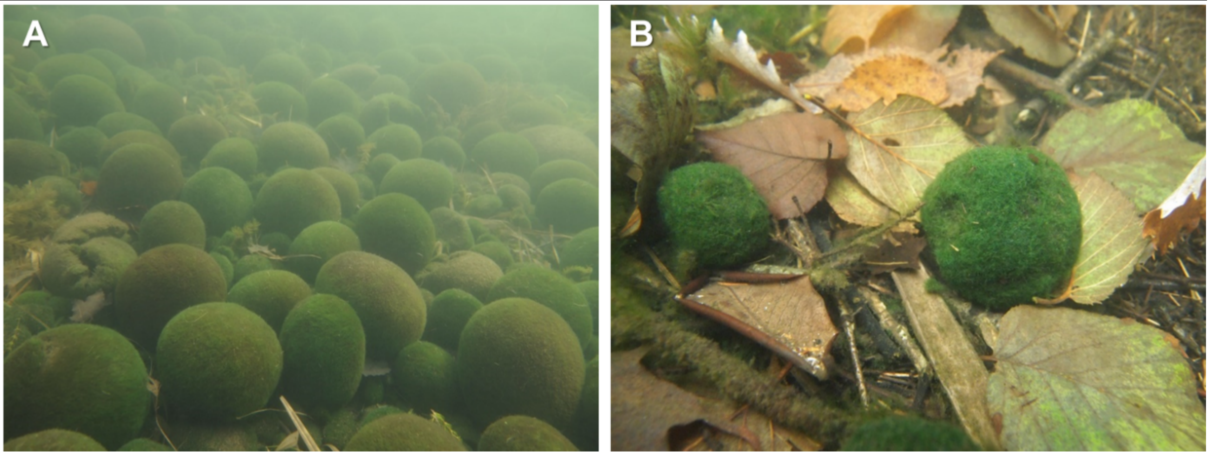


**Extended Data Figure 1. Marimo at the natural habitat of Lake Akan. (A)** Radial-type marimo are colonized on soft sediment. Water depth, 1.8 m. **(B)** Tangled-type marimo were scattered on the bottom of the lake, where leaves, pinecones, and wood branches were accumulated. Water depth, 0.5 m.

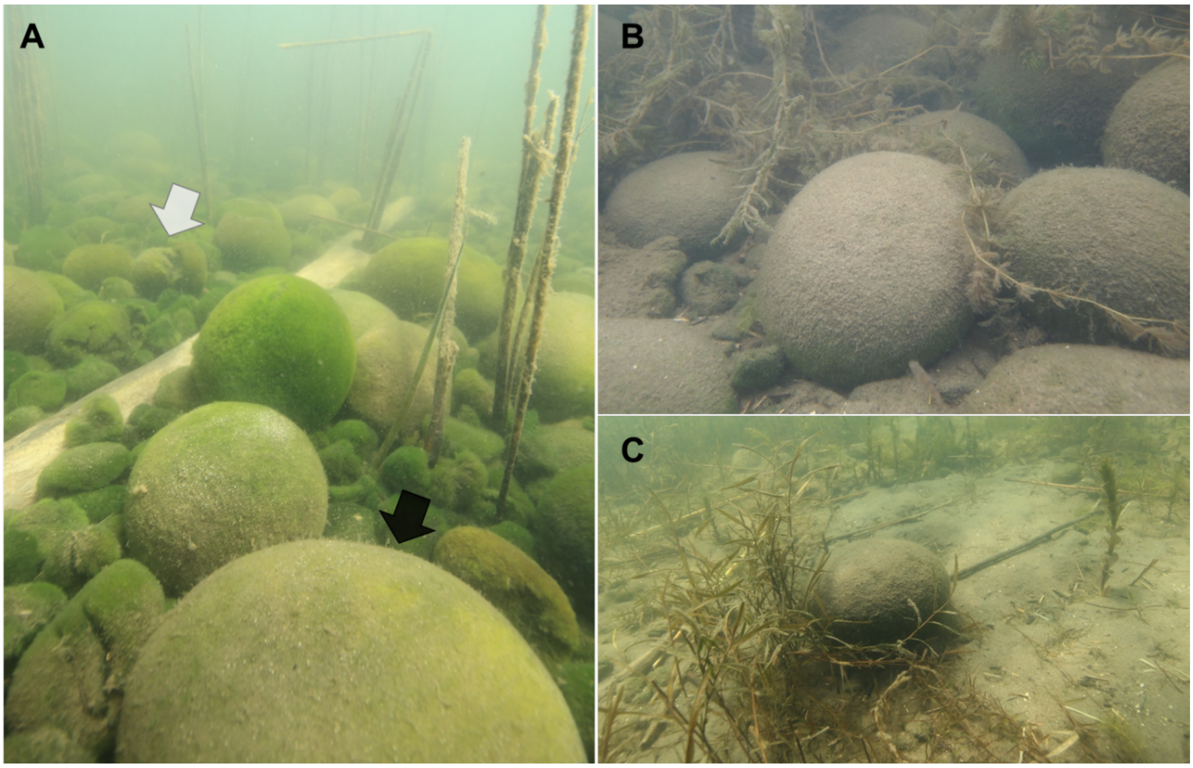


**Extended Data Figure 2. Images of the natural colonies of radial-type marimo distributed in a shallow region (water depth = 1.5 m) of Lake Akan. (A)** Marimo cover almost the entire lake bottom at the colony site. Some of the marimo are broken (*white arrow*). The marimo were slightly covered with lake sediments, which changed their surface color from green to brown (*black arrow*). **(B and C)** Marimo wholly covered by lake sediments. Sediment particles are removed by the rotatory motion of marimo generated by water current.

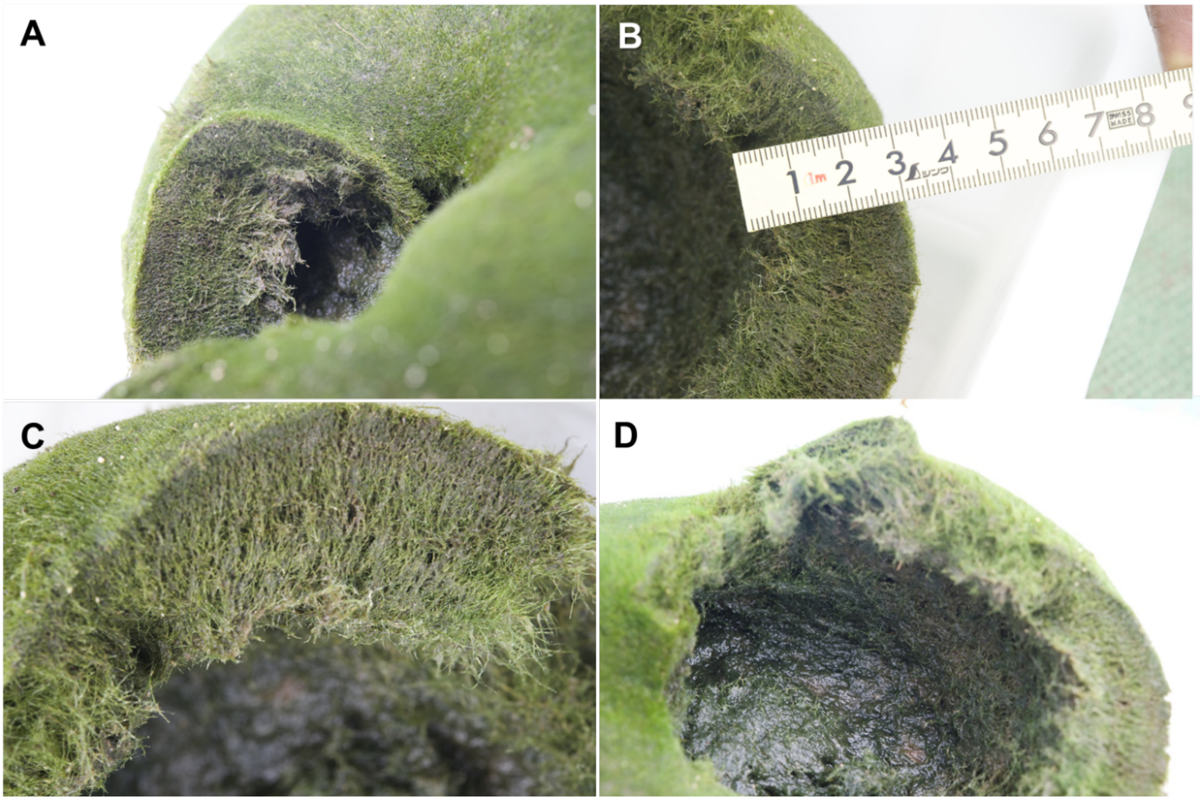


**Extended Data Figure 3. Images of cross sections of an algal mesh comprising a large radial-type marimo. (A to C)** Internal multi-layers in an algal outer mass (about 2.5 cm thick) contain brown-coloured sediment particles. See **Figure 3** for further details of the multi-layers. **(D)** A hollow structure formed in a large radial-type marimo.

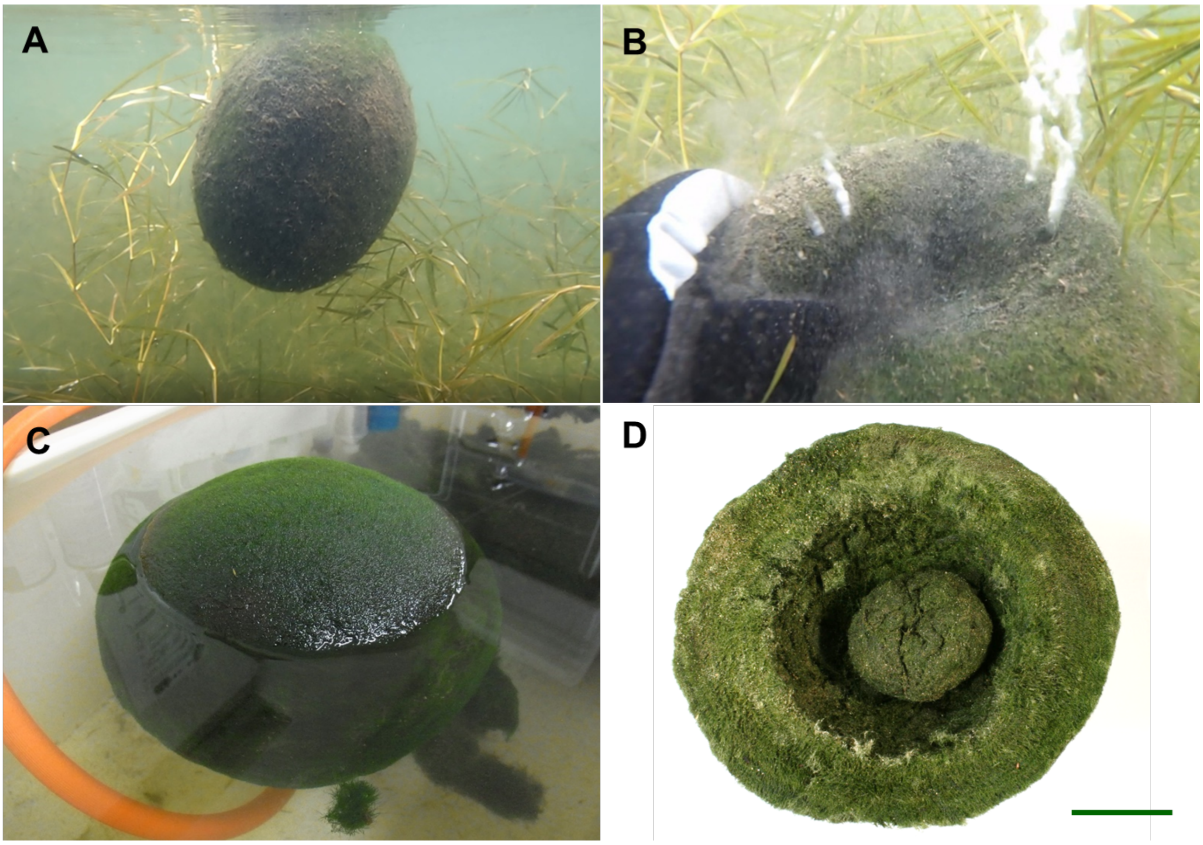


**Extended Data Figure 4. Sizable radial-type marimo with a large hollow structure. (A)** A buoyant radial-type marimo underneath the surface of Lake Akan. See **Supplementary Movie 1** for footage of the marimo in motion. **(B)** Gas extrudes from the internal hollow of the radial-type marimo after squeezing. **(C)** A buoyant radial-type marimo when internal water is completely removed after sampling from the lake bottom. **(D)** Image of the hollow structure developed only in large radial-type marimo after the removal of internal water. The individual contains a small aggregate regenerated from a marimo fragment peeled off inside the hollow. Scale bar, 5 cm.

**Supplementary Table 1** Sample codes and descriptions

| Code | Description |
|---|---|
| Radi-SO | outer surface section of the small radial-type marimo samples (about 4 cm in diameter) |
| Radi-SI | inner section of the small radial-type marimo samples (about 4 cm in diameter) |
| Radi-MO | outer surface section of the medium radial-type marimo samples (about 11 to 12 cm in diameter) |
| Radi-MI | inner section of the medium radial-type marimo samples (about 11 to 12 cm in diameter) |
| Radi-LO | outer surface section of the large radial-type marimo samples (about 21 and 22 cm in diameter) |
| Radi-LI | inner section of the large radial-type marimo samples (about 21 and 22 cm in diameter) |
| Fil | floating *Aegagropila linnaei* filaments |
| Tang-O | outer section of the tangled-type marimo samples (about 5 cm in diameter) |
| Tang-I | inner section of the tangled-type marimo samples (about 5 cm in diameter) |
| LW | surface lake water directly above the studied radial-type marimo colony |
| Sed-1 | lake-bottom sediment surrounding the studied small radial-type marimo samples |
| Sed-2 | lake-bottom sediment surrounding the studied medium radial-type marimo samples |
| Sed-3 | lake-bottom sediment surrounding the studied large radial-type marimo samples |
| Radi-L1 | brown-colored layer (~0.4 cm deep from the surface) formed in the large radial-type marimo samples |
| Radi-L2 | brown-colored layer (~1 cm deep from the surface) formed in the large radial-type marimo samples |
| Radi-L3 | brown-colored layer (~2 cm deep from the surface) formed in the large radial-type marimo samples |

**Supplementary Table 2** Bacteria-derived reads, OTU numbers, $\alpha$-diversity indices, and Good's coverages of amplicon data

| Sample code* | Target | Valid reads | # of OTUs | Chao1** | Non-parametric Shannon's index | Good's coverage of library (%) |
|---|---|---|---|---|---|---|
| Radi-SO | 16S rRNA gene | 143951 | 5865 | 5938.88 | 7.30 | 99.74 |
| Radi-SI | 16S rRNA gene | 151508 | 6485 | 6578.08 | 7.46 | 99.69 |
| Radi-MO | 16S rRNA gene | 177123 | 6244 | 6338.95 | 7.28 | 99.74 |
| Radi-MI | 16S rRNA gene | 108849 | 5028 | 5125.13 | 7.02 | 99.61 |
| Radi-LO | 16S rRNA gene | 209325 | 4664 | 4689.65 | 7.19 | 99.94 |
| Radi-LI | 16S rRNA gene | 186137 | 5997 | 6060.76 | 7.08 | 99.81 |
| Fil | 16S rRNA gene | 203967 | 4113 | 4171.31 | 6.55 | 99.86 |
| Tang-O | 16S rRNA gene | 144325 | 5612 | 5657.78 | 7.33 | 99.81 |
| Tang-I | 16S rRNA gene | 149830 | 4413 | 4477.78 | 6.46 | 99.79 |
| LW | 16S rRNA gene | 359695 | 4146 | 4389.90 | 4.46 | 99.80 |
| Sed-1 | 16S rRNA gene | 264429 | 8164 | 8336.97 | 6.90 | 99.71 |
| Sed-2 | 16S rRNA gene | 173522 | 7496 | 7668.29 | 7.11 | 99.56 |
| Sed-3 | 16S rRNA gene | 128709 | 4390 | 4476.00 | 6.54 | 99.70 |
| Radi-L1 | 16S rRNA transcript | 113725 | 5719 | 5751.37 | 7.42 | 99.79 |
| Radi-L2 | 16S rRNA transcript | 84188 | 4281 | 4306.68 | 7.08 | 99.78 |
| Radi-L3 | 16S rRNA transcript | 189170 | 6826 | 6850.89 | 7.30 | 99.87 |

* Detailed information on sample codes is given in **Table S1**.

** This index is calculated based on the number of the rare observed species (singleton and doubleton reads)

**Supplementary Table 3** Chemical composition and environmental parameters of lake surface water above the marimo colony studied in Churui bay, Lake Akan, Japan.

| Compound/parameter | Data value |
|---|---|
| pH | 8.01 |
| Electrical conductivity (ms $m^{-1}$) | 23.60 |
| $Na^+$ (mg $L^{-1}$) | 17.0 |
| $K^+$ (mg $L^{-1}$) | 2.95 |
| $Ca^{2+}$ (mg $L^{-1}$) | 15.0 |
| $Mg^{2+}$ (mg $L^{-1}$) | 7.26 |
| $NH_4^+$ (mg $L^{-1}$) | <0.05* |
| $Cl^-$ (mg $L^{-1}$) | 18.6 |
| $NO_2^-$ (mg $L^{-1}$) | <0.02* |
| $NO_3^-$ (mg $L^{-1}$) | <0.05* |
| $PO_4^{3-}$ (mg $L^{-1}$) | <0.10* |
| $SO_4^{2-}$ (mg $L^{-1}$) | 37.4 |
| Total nitrogen (mg $L^{-1}$) | 0.07 |
| Total phosphorus (mg $L^{-1}$) | <0.01* |
| Total organic carbon (mg $L^{-1}$) | 1.30 |

* below the detection limit

**A note on Supplementary Tables 4 to 9**

Please note that Supplementary Tables 4−9 listing amplicon read counts and their taxonomic classification at different taxonomic levels (phylum, class, order, family, genus, and species) are large files, therefore, summarizing Excel data are uploaded separately.

**Supplementary Table 4** Amplicon read counts and taxonomic classification derived from the studied marimo samples and comparative samples at the phylum level

**Supplementary Table 5** Amplicon read counts and taxonomic classification derived from the studied marimo samples and comparative samples at the class level

**Supplementary Table 6** Amplicon read counts and taxonomic classification derived from the studied marimo samples and comparative samples at the order level

**Supplementary Table 7** Amplicon read counts and taxonomic classification derived from the studied marimo samples and comparative samples at the family level

**Supplementary Table 8** Amplicon read counts and taxonomic classification derived from the studied marimo samples and comparative samples at the genus level

**Supplementary Table 9** Amplicon read counts and taxonomic classification derived from the studied marimo samples and comparative samples at the species level

**A note on Supplementary Movie 1**

Please note that Supplementary Movie 1 shows a buoyant radial-type marimo underneath the surface of Lake Akan.
